# Chemotaxonomic Investigation of Apocynaceae for Retronecine-Type Pyrrolizidine Alkaloids Using HPLC-MS/MS

**DOI:** 10.1101/2020.08.23.260091

**Authors:** Lea A. Barny, Julia A. Tasca, Hugo A. Sanchez, Chelsea R. Smith, Suzanne Koptur, Tatyana Livshultz, Kevin P. C. Minbiole

## Abstract

Apocynaceae are well-known for diverse specialized metabolites that are distributed in a phylogenetically informative manner. Pyrrolizidine alkaloids (PAs) have been reported sporadically in one lineage in the family, the APSA clade, but few species had been studied to date. We conduct the first systematic survey of Apocynaceae for retronecine-type PAs, sampling leaves from 231 species from 13 of 16 major lineages within the APSA clade using HPLC-MS/MS. We also follow up on preliminary evidence for infra-specific variation of PA detectability in *Echites umbellatus* Jacq. Four precursor ion scans (PREC) were developed for a high-throughput survey for chemicals containing a structural moiety common to many PAs, the retronecine core. We identified with high confidence PAs in 7 of 8 sampled genera of tribe Echiteae, but not in samples from the closely related Odontadenieae and Mesechiteae, confirming the utility of PAs as a taxonomic character in tribal delimitation. The presence of PAs in Malouetieae was confirmed, as we report with high confidence their presence in *Galactophora schomburgkiana* Woodson and *Eucorymbia alba* Stapf, but currently we have low confidence of their presence in *Holarrena pubescens* Wall. ex G. Don (the one Malouetieae species where they were previously reported), as well as in *Kibatalia macrophylla* (Pierre ex Hua) Woodson and in *Holarrena curtisii* King & Gamble. For the first time the presence of PAs in species of *Wrightia* R. Br. (Wrightieae) and *Marsdenia* R. Br. (Marsdenieae) was confirmed. Detectability of PAs was found to vary among samples of *Echites umbellatus* and intra-individual plasticity contributes to this variation. Of toxicological importance, novel potential sources of human exposure to pro-toxic PAs were identified in the medicinal plants, *Wrightia tinctoria* R.Br. and *Marsdenia tinctoria* R.Br., and the food plant, *Echites panduratus* A. DC., warranting immediate further research to elucidate the structures of the candidate PAs identified. Method development and limitations are discussed.

## 1. Introduction

The study of plant specialized metabolites in a phylogenomic framework promises to fulfill the original goal of chemotaxonomy: identification of lineage diagnostic metabolites (Fairbrothers et al., 1975). In combination with other –omic, biochemical, and ecological approaches, chemotaxonomy permits the testing of key hypotheses about the evolutionary assembly and fates of novel metabolic pathways (Scossa and Fernie, 2020) as well as the role of plants’ interactions with their environments, including co-evolution with their specialized herbivores, in shaping plant metabolic diversity (Futuyma and Agrawal, 2009). One of the primary impediments to pursuit of this research agenda in large plant families is the fragmentary knowledge of the taxonomic occurrence of specialized metabolites. The published literature on natural products contains multiple biases, with the absence of a class of metabolites in a species being particularly hard to infer from studies that have not specifically targeted a compound class. Systematic surveys for chemicals of interest are often also impeded by availability of tissues across a breadth of species; this can be compounded by the lack of high-throughput, sensitive and selective analytical methods to maximize compound identification from small quantities of test material (Tasca et al., 2018). Infraspecific variation, a ubiquitous feature of plant specialized metabolism (Hartmann, 1996; Moore et al., 2014), can further introduce confusion about the distribution of a metabolite via contradictory reports in the literature.

Apocynaceae are the tenth largest angiosperm plant family, with ca. 5300 species classified in 378 genera (Endress et al., 2018). The family is of particular importance in natural products research because of the occurrence of multiple medicinally important species and compounds including monoterpenoid indole alkaloids, particularly the chemotherapy drugs vincristine and vinblastine in *Catharanthus roseus* (L.) G.Don (Aslam et al., 2010). The expense and difficulty of deriving these valuable molecules from natural sources has motivated the complete biochemical elucidation of their biosynthetic pathway as a steppingstone to its genetic engineering (Caputi et al., 2018; Qu et al., 2019). The genus *Asclepias* L. and its specialized herbivores are a model system in chemical ecology and evolution of reciprocal adaptations between plants and herbivores, with a particular focus on cardenolides (Agrawal et al., 2012). Pyrrolizidine alkaloids are also implicated in co-evolution between Apocynaceae and one of their specialized herbivore lineages, Lepidoptera subfamily Danainae (milkweed and clearwing butterflies), and the evolution of the first gene of their biosynthetic pathway, homospermidine synthase (*hss*), has been elucidated (Livshultz et al., 2018a). Researchers have detected phylogenetic signals in the distribution of all of these compounds and others, including steroidal and phenanthroindolizidine alkaloids and steroidal glycosides, in Apocynaceae (Endress et al., 2018), but knowledge of their taxonomic distribution still lags progress on the phylogeny of the family (Fishbein et al., 2018).

**Pyrrolizidine alkaloids** (PAs) are secondary metabolites containing a 1-hydroxymethylated necine core, esterified with a necic acid. The necine base (often referred to as a pyrrolizidine) presents as a heterocycle containing a nitrogen atom positioned at the junction of two fused five-membered rings (Hartmann and Witte, 1995) (Figure 1). Depending on the necine base present, PAs can be classified into five main types: retronecine (155 Da) or its less common diastereomer, heliotridine (155 Da), the monocyclic otonecine (185 Da) (These et al., 2013), the fully saturated platynecine (157 Da) (Lin et al., 1998), or the triol rosmarinecine (173 Da) (De Waal, 1941).

**Fig. 1.**
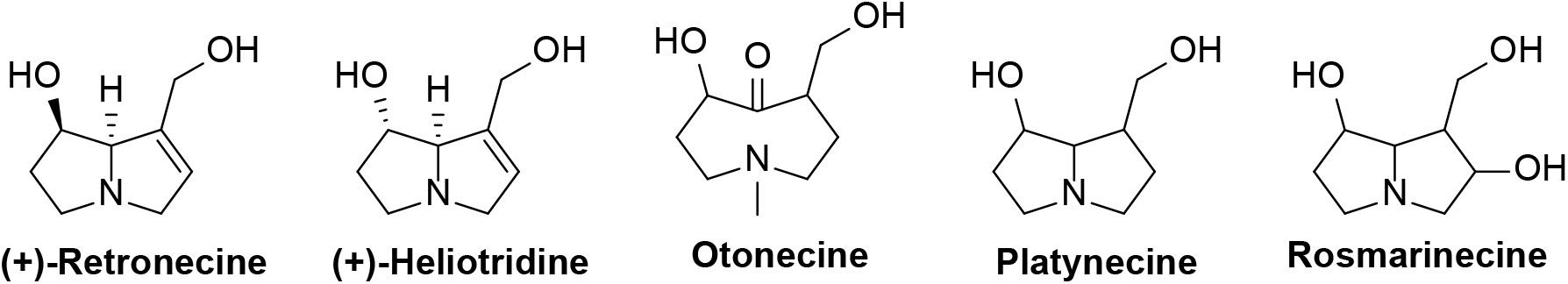
Necine bases. Varied necine base structures found in pyrrolizidine alkaloids.

Additionally, due to the immense structural diversity imparted by various combinations of necine bases with necic acids, PAs are primarily characterized according to esterification pattern and necic acid type (Hartmann and Witte, 1995) (Fig. 2).

**Fig. 2.**
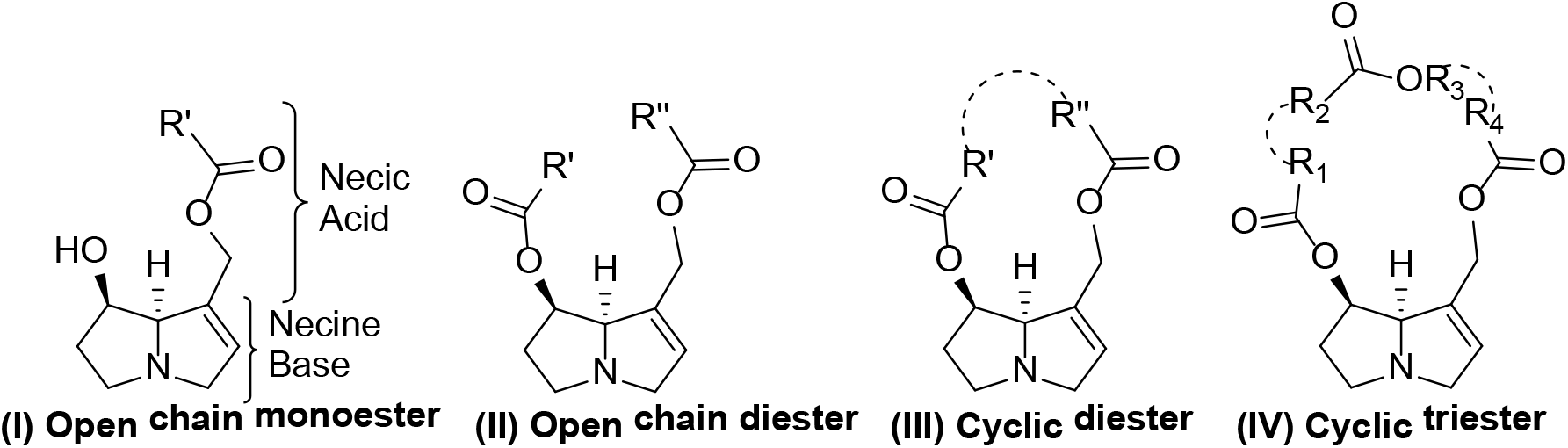
PA esterification patterns. Common esterification patterns between necic acids and necine bases observed in retronecine-type PAs.

PAs are known in 12 plant families (Hartmann and Witte, 1995; Tamariz et al., 2018) in addition to Apocynaceae. Previous studies exploring the structural diversity of PAs within Apocynaceae have primarily noted the presence of lycopsamine-type PAs, group abbreviation “C” (Hartmann and Witte, 1995), containing various subgroups: lycopsamine, isolycopsamine, latifoline, and parsonsine, referred to as C1, C2, C3, and C4, respectively (Burzynski et al., 2015; Colegate et al., 2016) (Fig. 3). Various other types of PAs are also present within Apocynaceae, including phalaenopsine (E) and miscellaneous (M; comprised of unusual necine esters/ simple necine derivates) (Burzynski et al., 2015). Other PA classes, senecionine (referred to as A), triangularine (B), monocrotaline (D), and loline (L), have never been reported from Apocynaceae (Hartmann and Witte, 1995).

**Fig. 3.**
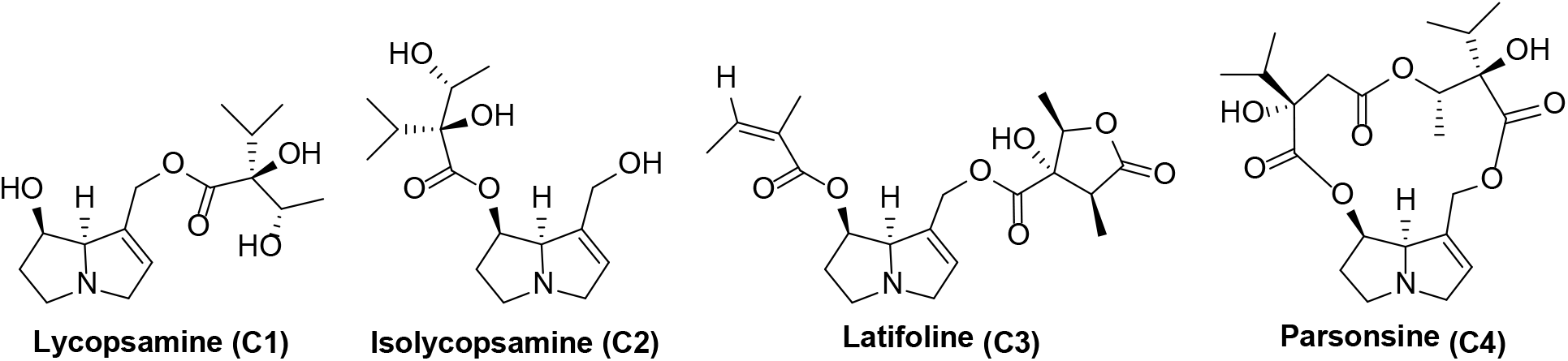
Lycopsamine-type PAs. Subtypes of lycopsamine-type PAs containing a retronecine core.

The chemical ecology of PAs is among the most intensely studied of any group of plant specialized metabolites, with multiple origins of PA-phily among diverse insect lineages (Hartmann and Witte, 1995; Trigo, 2011). PAs are important in agriculture because of their toxicity to livestock (Lucena et al., 2010), and as biocontrol against phytophagous nematodes (Thoden and Boppré, 2010). Their presence in foods and herbal preparations is of concern to public health regulators, given the toxicity of desaturated PAs (Dusemund et al., 2018; Kaltner et al., 2020; Schramm et al., 2019).

To date, only 37 Apocynaceae species have been specifically tested for PAs via chemical means, with 15 PA-positive species discovered in 7 genera (reviewed in Burzynski et al, 2015; Colegate at al., 2016). These genera belong to four distantly related tribes: Nerieae (*Alafia* Thouars), Malouetieae (*Holarrhena* R.Br.), Echiteae (*Echites* P.Browne*, Prestonia* R.Br., *Parsonsia* R.Br.), Apocyneae (*Anodendron* A.DC., *Amphineurion* (A.DC.) Pichon). Report of PAs in *Macropharynx* Rusby (syn. *Peltastes* Woodson), *Temnadenia* Miers, and *Thenardia* Kunth (Echiteae) (Morales et al., 2017) are derived from observed behaviors of PA-philous insects rather than chemical analysis (Brown, 1987; Hernández-Baz et al., 2013). Based on the evolution of *hss*-like genes in Apocynaceae, Livshultz et al. (2018a) proposed that PAs evolved in the common ancestor of these four tribes, early in the diversification of a well-corroborated lineage, the APSA clade, followed by multiple independent losses. This implies that PAs may be more widely distributed within the APSA clade, which includes ca. 4000 species, than reported in the literature. Conflicting reports of presence or absence in a particular species, such as *Prestonia coalita* (Vell.) Woodson (Burzynski et al., 2015), and variation in PA detectability among herbarium specimens of *Echites umbellatus* Jacq. (Tasca et al., 2018), indicate the potential for infra-specific variation in PA presence.

### 1.2. Scanning for Retronecine-Type PAs via HPLC-MS/MS

The abundance of compounds classified as PAs, greater than 400 to date, and the immense structural diversity exhibited by PAs, including both isobaric constitutional and stereoisomers (Hartmann and Witte, 1995), in conjunction with a relative lack of available standards, has led to the advancement of broadly useful PA detection methods (Jeong and Lim, 2019; These et al., 2013). These techniques do not require corresponding standards as they target key structural fragments of PAs by subjecting them to mass spectrometry (MS) coupled to either liquid or gas chromatography (Jeong and Lim, 2019; Stelljes et al., 1991). In the case of retronecine/heliotridine-type PAs, previous groups have utilized the m/z 120 and 138 fragment ions to detect cyclic diesters and open-chained diesters (Figure 4). Since retronecine and heliotridine -type PAs only differ by their stereochemical configuration, they present with the same structurally significant fragments upon MS analysis (These et al., 2013). Additionally, m/z 120, 138, and 156 fragment ions have been used to identify open-chained monoesters (Avula et al., 2015; Avula, 2015; Lin et al., 1998; Sixto et al., 2019; These et al., 2013) (Figure 4). Fragments of the retronecine/heliotridine core have also been used as qualifier ions in PA identification via mass spectrometry. The major sub-fragments of the retronecine core include m/z 94 and m/z 96, with m/z 94 being more common in PA characterization (Colegate et al., 2016; Lin et al., 1998; Pedersen and Larsen, 1970).

**Fig. 4.**
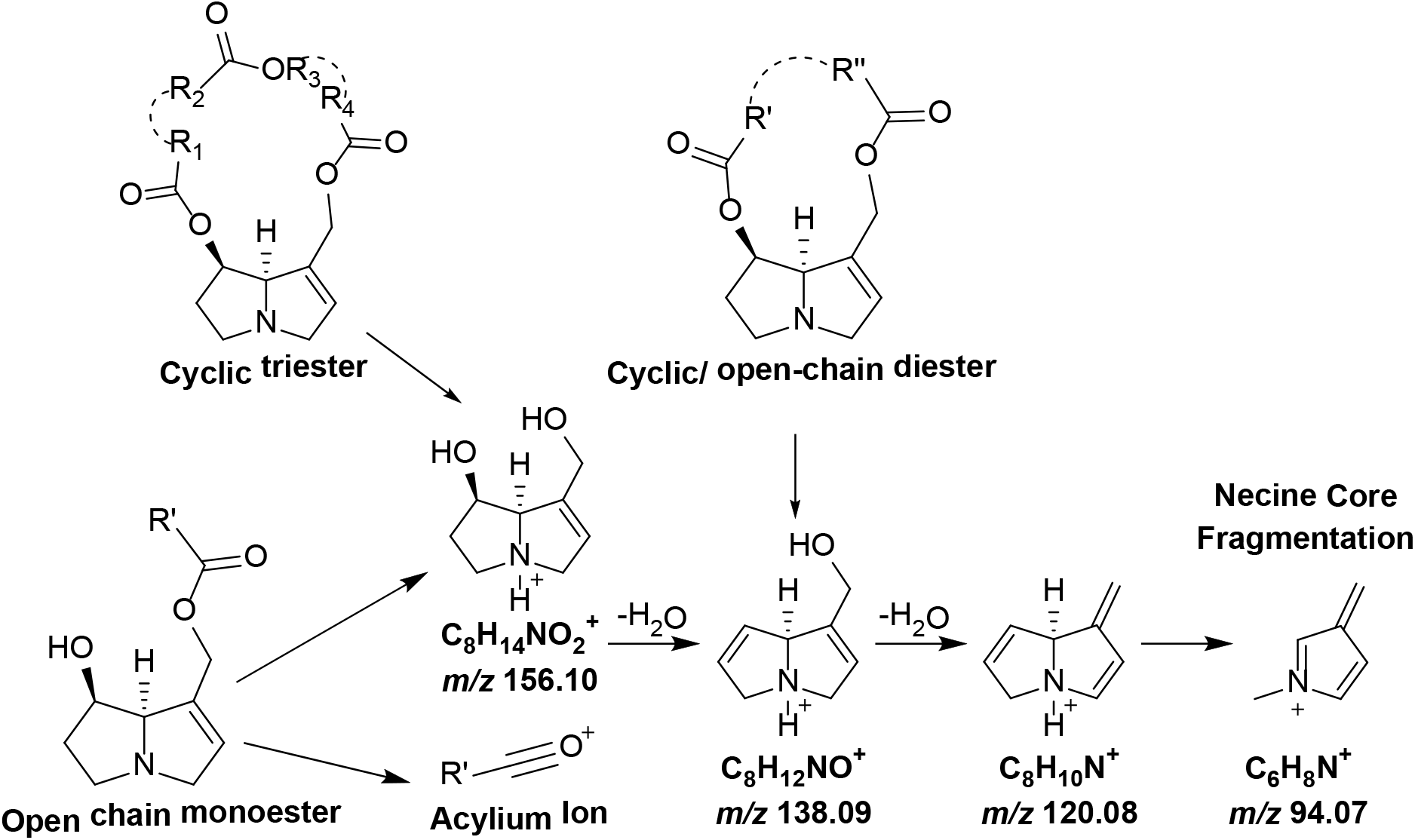
PA Fragmentation scheme. Fragmentation patterns according to esterification pattern in retronecine-type PAs.

Burzynski et al. (2015) used multiple types of MS scans (Q1, MRM, PREC) to tentatively identify specific PAs, as both free bases and N-oxides, known to be part of the parsonsine (C4) subgroup of lycopsamine-type PAs in Apocynaceae species (Hartmann and Witte, 1995). MRM (multiple reaction monitoring) and PREC (precursor ion) scans were generated to identify the m/z 120 fragment, typical of retronecine-type PAs. Leaf, seed, nectar, and sap samples of *Echites umbellatus* and *Parsonsia alboflavescens* (Dennst.) Mab. (Echiteae) were all found to contain PAs with parsonsine isomers likely present.

Alternatively, to determine total PA content in honey samples, Sixto et al. (2019) utilized MS precursor, product, and multi-reaction monitoring (MRM) scans, similar in experimental design to These et al. (2013). Precursor scans were designed to locate evenly massed molecular ions [M+H]^+^ with either m/z 120 (open chain mono/di-esters, cyclic diesters, or cyclic diester N-oxides) or m/z 138 fragments (monoester N-oxides). Once molecular ion precursors associated with retronecine/heliotridine-type Pas were established, these molecular ions were then subject to product scans to evaluate overall fragments, with MRMs being utilized for quantification.

In the current study, to survey for the presence of retronecine/heliotridine-type PAs and avoid the need to generate multiple product scans for the proposed PA molecular ions, PREC scans for the major fragments of retronecine/heliotridine-type PAs were used to identify potential PA molecular ions in positive ion scanning mode. Since nearly all PAs contain a single nitrogen atom within their pyrrolizidine ring, and because we exploited positive electrospray ionization, we selectively targeted species with even mass to charge (m/z) ratio. MCA (multichannel analysis) was used in PREC 120 scans for initial sample screening. This function produced an additive spectrum with increased sensitivity, reducing the chance of falsely identifying samples as negative. However, MCA was not used when samples were subject to a secondary analysis, involving the use of three PREC scans (m/z 120, 138, and 156) to allow for evaluation of the mass spectrum throughout the entirety of the chromatographic run. Our LC-MS methods were validated on a set of purchased PA standards, as well as on positive and negative control species.

We report the largest survey to date of PA occurrence in Apocynaceae, leveraging the long-term stability of PAs in dried leaves (Tasca et al., 2018), the taxonomically diverse collection of leaf samples that one of us (T.L.) has assembled for phylogenetic analysis, and advances in PA identification using HPLC-MS/MS to overcome the barriers that had previously precluded systematic study of their occurrence. We also follow up on preliminary evidence of infraspecific polymorphism of PA presence in *Echites umbellatus* (Tasca et al., 2018) by increasing infraspecific sampling and re-sampling individual plants over time to infer the presence of intra-individual plasticity. Finally, we apply our methods to the analysis of seeds of two medicinal species: *Holarrhena pubescens* Wall. & G.Don and *Wrightia tinctoria* R.Br.

## 2. Results and discussion

### 2.1. Method validation

All purchased PA standards (heliotrine, jacobine, retrorsine, europine, lycopsamine and monocrotaline), representing cyclic diesters and open-chained monoesters, resulted in chromatographic separation in the TIC (total ion chromatograph) of PREC 120 and 138 scans. Scans of the open-chained monoesters (europine, lycopsamine, and heliotrine) also resulted in distinct features in the TIC of the PREC 156 scan. All six standards appeared in the TIC (total ion chromatogram) between 9-12 minutes (45-60% ACN), using the chromatographic conditions outlined below.

*Apocynum cannabinum* L. was used as a negative control because it is a chemically well-studied species from which PAs have not been reported (Abe and Yamauchi, 1994), and because it has a homospermidine synthase pseudogene and hence should not be able to produce PAs (unpublished data). Subjecting the negative control leaf sample to the four PREC scans resulted in a TIC with a signal to noise ratio lower than three to one (<3:1 S/N).

*Parsonsia alboflavescens* served as the positive control for method validation of the PREC scans because its PAs have been previously characterized by NMR/ MS (Colegate et al., 2016). We identified 18 candidate PAs in *P. alboflavescens* (Table 1) by targeting common structural fragments of retronecine-type PAs via multiple PREC scans (m/z 120, 138, and 156). Fourteen (14) of these are identical in mass to PAs previously identified in *Parsonsia* spp. via NMR (Abe et al., 1991a; Abe et al., 1990; Abe and Yamauchi, 1987; Abe et al., 1991b; Edgar et al., 1980; Eggers and Gainsford, 1979; Nishida et al., 1991), and 11 match masses previously identified by PREC 120 (Burzynski et al., 2015). However, without HRMS analysis and NMR structural elucidation, which is difficult on the sample scale utilized, molecular ion structural determinations and identifications remain tentative.

**Table 1.**
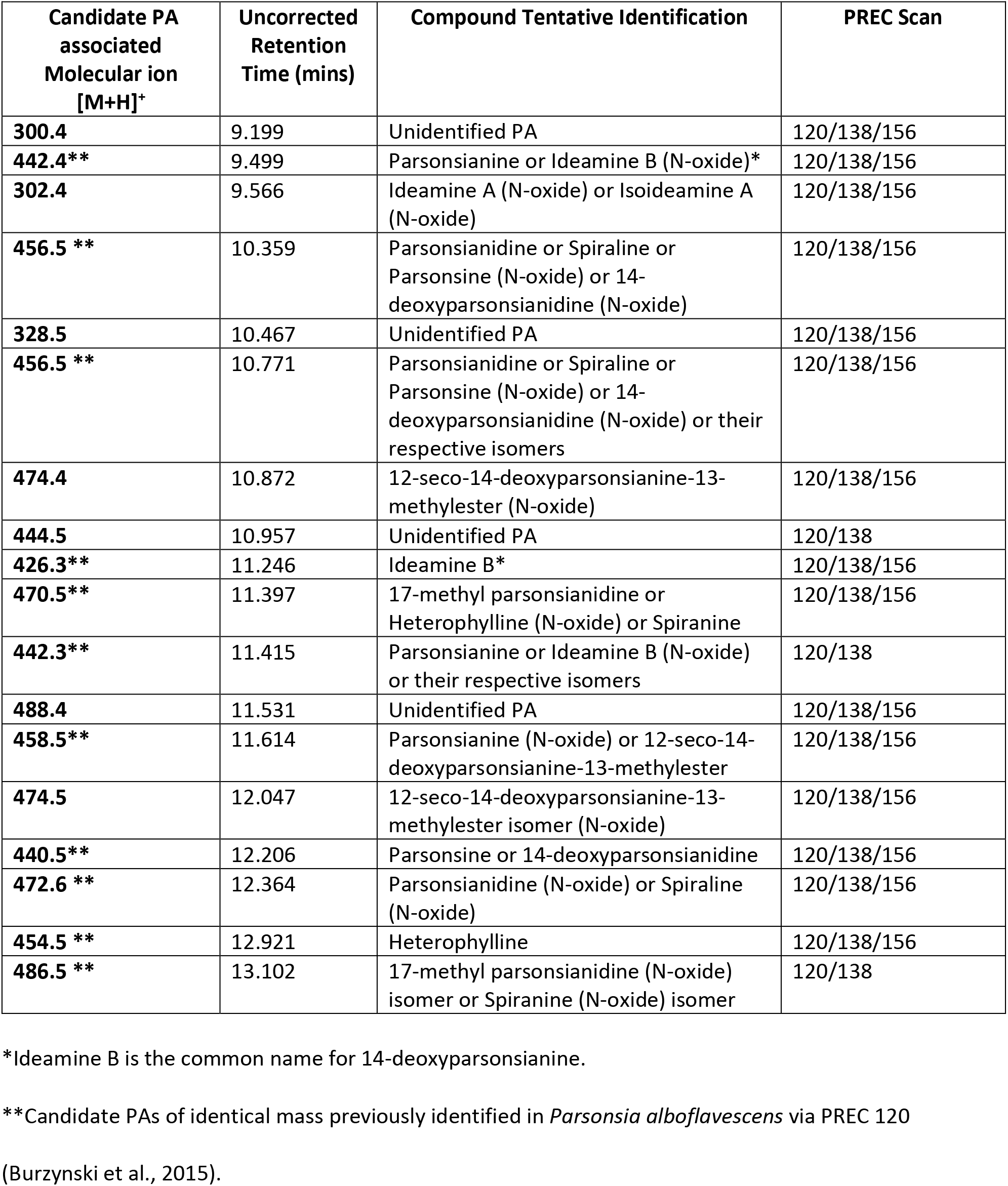
Candidate PAs identified in *Parsonsia alboflavescens*. Voucher in Table S1.

| Candidate PA associated Molecular ion [M+H]<sup>+</sup> | Uncorrected Retention Time (mins) | Compound Tentative Identification | PREC Scan |
| --- | --- | --- | --- |
| 300.4 | 9.199 | Unidentified PA | 120/138/156 |
| 442.4** | 9.499 | Parsonsianine or Ideamine B (N-oxide)* | 120/138/156 |
| 302.4 | 9.566 | Ideamine A (N-oxide) or Isoideamine A (N-oxide) | 120/138/156 |
| 456.5 ** | 10.359 | Parsonsianidine or Spiraline or Parsonsine (N-oxide) or 14-deoxyparsonsianidine (N-oxide) | 120/138/156 |
| 328.5 | 10.467 | Unidentified PA | 120/138/156 |
| 456.5 ** | 10.771 | Parsonsianidine or Spiraline or Parsonsine (N-oxide) or 14-deoxyparsonsianidine (N-oxide) or their respective isomers | 120/138/156 |
| 474.4 | 10.872 | 12-seco-14-deoxyparsonsianine-13-methylester (N-oxide) | 120/138/156 |
| 444.5 | 10.957 | Unidentified PA | 120/138 |
| 426.3** | 11.246 | Ideamine B* | 120/138/156 |
| 470.5** | 11.397 | 17-methyl parsonsianidine or Heterophylline (N-oxide) or Spiranine | 120/138/156 |
| 442.3** | 11.415 | Parsonsianine or Ideamine B (N-oxide) or their respective isomers | 120/138 |
| 488.4 | 11.531 | Unidentified PA | 120/138/156 |
| 458.5** | 11.614 | Parsonsianine (N-oxide) or 12-seco-14-deoxyparsonsianine-13-methylester | 120/138/156 |
| 474.5 | 12.047 | 12-seco-14-deoxyparsonsianine-13-methylester isomer (N-oxide) | 120/138/156 |
| 440.5** | 12.206 | Parsonsine or 14-deoxyparsonsianidine | 120/138/156 |
| 472.6 ** | 12.364 | Parsonsianidine (N-oxide) or Spiraline (N-oxide) | 120/138/156 |
| 454.5 ** | 12.921 | Heterophylline | 120/138/156 |
| 486.5 ** | 13.102 | 17-methyl parsonsianidine (N-oxide) isomer or Spiranine (N-oxide) isomer | 120/138 |

\*Ideamine B is the common name for 14-deoxyparsonsianine.
\*\*Candidate PAs of identical mass previously identified in *Parsonsia alboflavescens* via PREC 120
(Burzynski et al., 2015).

At the declustering potentials and collision energies utilized by the mass spectrometer, [M+H]^+^ ions corresponding to the masses of predicted cyclic triesters largely produced all three fragments characteristic of retronecine-type PAs, m/z 120, 138, and 156. While retronecine-type PAs are structurally similar, with fragments understood to involve the necine base and the hydroxymethyl moiety positioned at carbon one of the pyrrolizidine ring, the subtle differences in chemical structures and analytical instrumentation prevents absolute optimization of collision energy and declustering potential values utilized in PREC scans; these instrumental parameters are still largely compound dependent, and therefore can result in inadequate sensitivity for the detection of a particular fragment, resulting in the possibility of false negatives.

### 2.2. Survey for PA distribution

A total of 319 samples from 231 species [representing 16 of 28 major Apocynaceae lineages classified as subfamilies or tribes (Endress et al., 2018)] were screened for PAs utilizing a PREC 120 scan with MCA (vouchers and results in the supporting Information Table S1). Samples containing a molecular ion with a m/z 120 fragment were determined to be positive, with low confidence, for retronecine-type PAs based on following criteria: (1) the TIC (total ion chromatogram) signal, associated with an evenly massed molecular ion, was greater than 3:1 S/N with column retention between 9 and 15 minutes, and (2) the evenly massed molecular ion of interest was greater than or equal to 1×10^4^ intensity (cps; counts per second). Select samples were then evaluated on PREC 120/ 138/ 156 methods to confirm additional fragments of evenly massed molecular ions suspected to be PAs (Table 2).

**Table 2.**
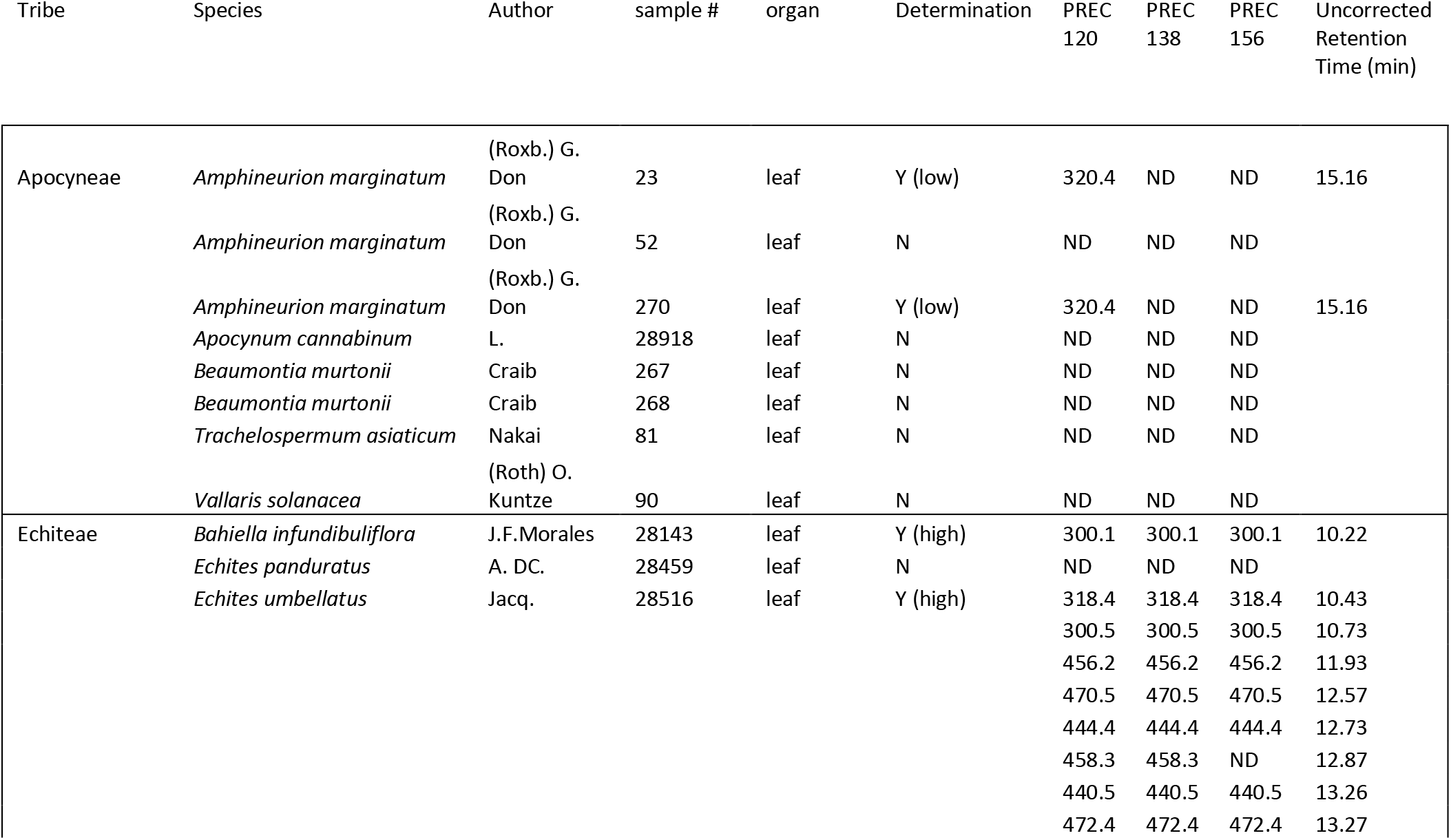
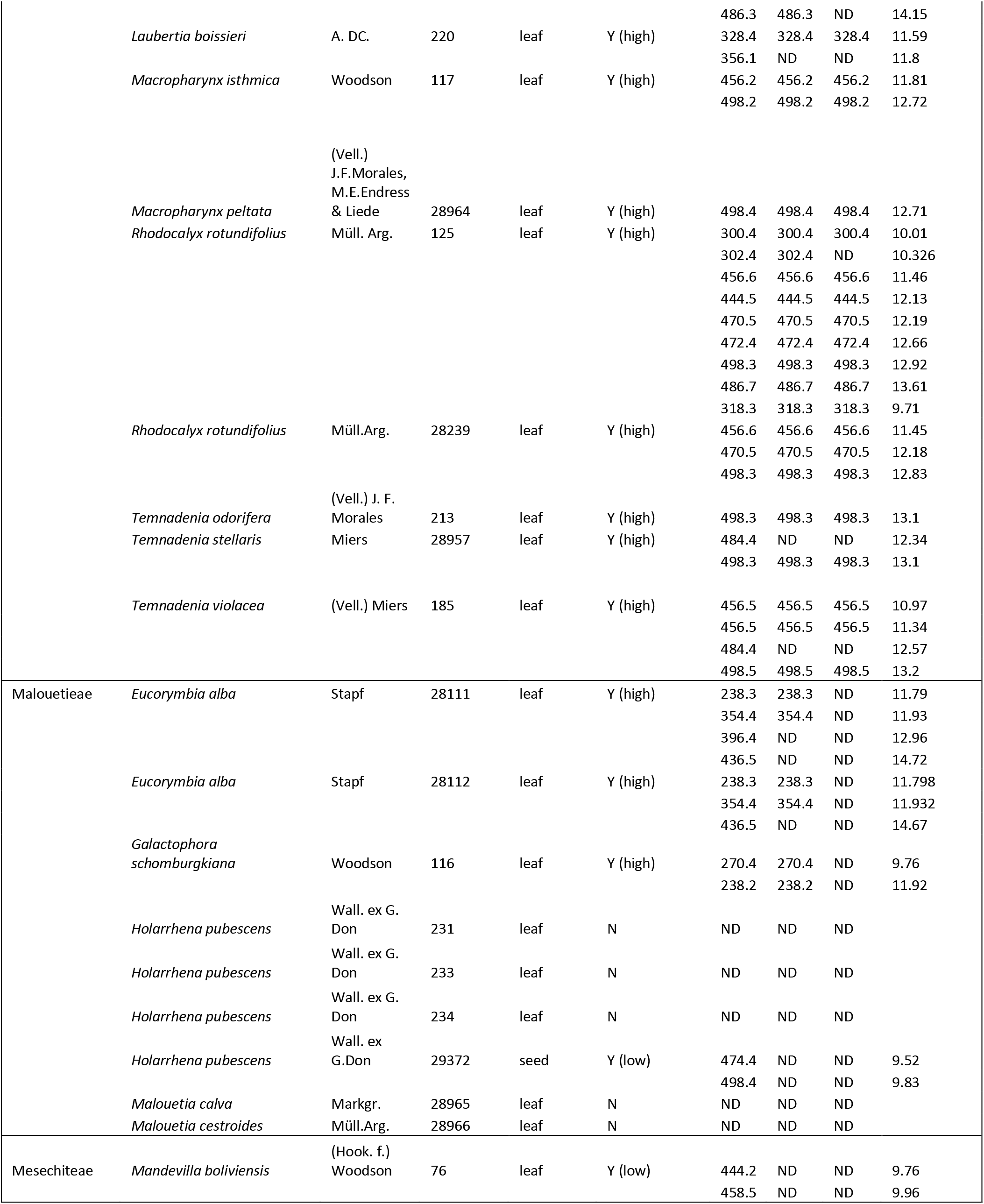
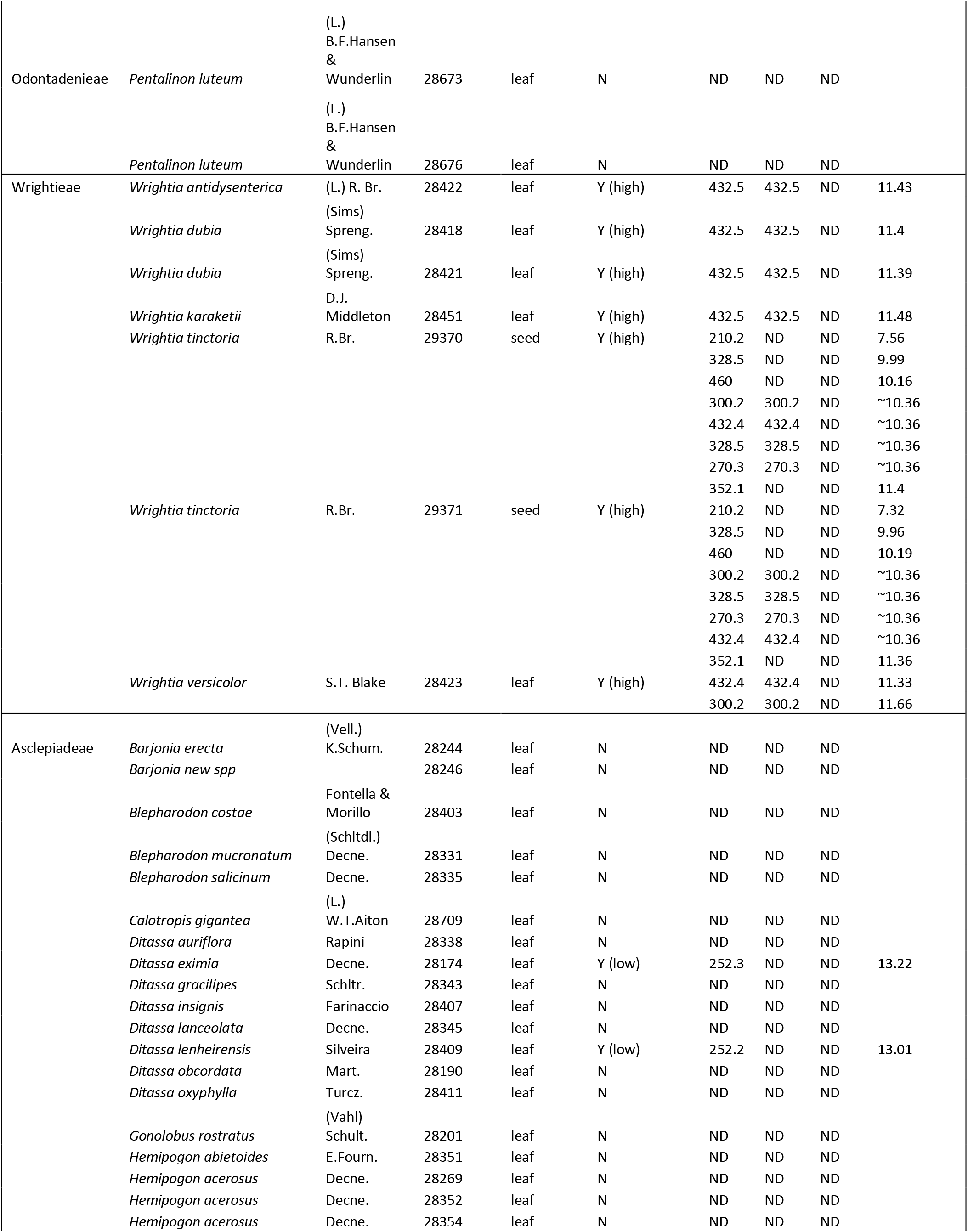
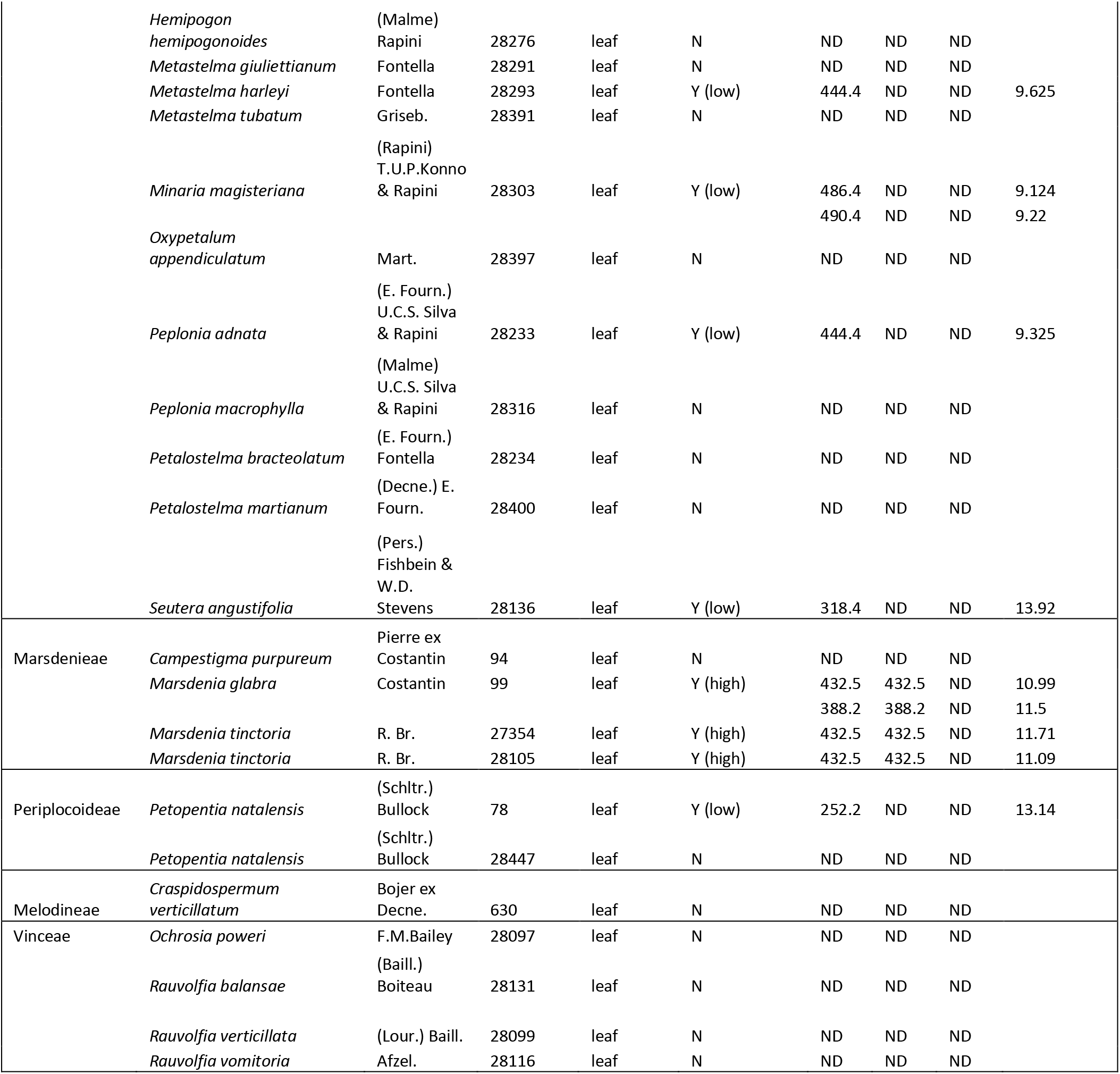
Samples analyzed on all PREC scans (m/z 120, 138, 156). Samples determined as N: no candidate ions detected, Y (low): low confidence, candidate ions detected only in PREC 120, and Y (high): high confidence, candidate ions detected in PREC 120 and PREC 138. ND: not detected. Vouchers in Table S1.

Samples were deemed as “positive with high confidence” if both the PREC 120 and PREC 138 scans indicated the same ion at the same retention time. As previously indicated, molecular ions that fragment to m/z 120 and 138 are typically associated with cyclic- diesters and open- chain diesters, and PA compounds whose fragments include m/z 120, 138, and 156 are likely to be open-chained monoesters and cyclic triesters (Fig. 4). Samples that showed no strong features on PREC 138 or 156 scans were deemed “positive with low confidence” for retronecine-type PAs. Of the 78 leaf samples analyzed via all PREC methods (m/z 120, 138, 156; without MCA), 45 were determined to be negative or contain PAs below the limit of detection of the method (Table 2). Ten (10) samples from species of Apocyneae, Asclepiadeae, Periplocoideae, and *Holarrhena pubescens* (Malouetieae) were “positive with low confidence” since they yielded only ions with an m/z 120 fragment. Of the “positive with high confidence” samples (23), those from species of Malouetieae, Wrightieae, and Marsdenieae contained likely PAs with only m/z 120 and 138 fragments, while all positive samples from Echiteae species contained one or more candidate PAs with m/z 120, 138, and 156 fragments.

### 2.3. Phylogenetic distribution of PAs

None of the Rauvolfioid taxa sampled (11 species from 5 genera in 3 of 11 tribes, Tabernaemontaneae, Melodineae, and Vinceae) had any detectable PAs (Table S1, Table 2). These taxa lack a homospermidine synthase-like gene (Livshultz et al., 2018a) and are therefore predicted to lack the capacity for PA biosynthesis. They are known for the production of monoterpenoid indole alkaloids (MIAs) (Endress et al., 1990), which our methods would be unlikely to detect.

All ions identified as PAs with high confidence (Table 2) occur in species of tribes of the APSA clade: two previously reported (Echiteae, Malouetieae) (Burzynski et al., 2015) and two newly reported here (Wrightieae, Marsdenieae). Ions identified as PAs with low confidence were detected in species of tribes Echiteae, Malouetieae, and Apocyneae (*Amphineurion*, previously reported) (Colegate et al., 2016) and Wrightieae, Asclepiadeae, Mesechiteae, and Periplocoideae (newly reported). None of seven sampled species from the five (of six total) genera in tribe Nerieae had detectable ions in the PREC 120 (with MCA) scans (Table S1). *Alafia* cf. *caudata* Stapf (Nerieae) had been previously reported to produce retronecine-type PAs (Colegate et al., 2016), but we did not detect any candidate compounds in samples of *Alafia barteri* Oliv. or *A. thouarsii* Roem. & Schult. *Alafia* includes 22 species (Leeuwenberg, 1997; Wieringa, 2011) and further sampling may identify chemotaxonomically useful variation in PA profiles. All samples from other major APSA clade lineages (Rhabdadenieae, Odontadenieae, Baisseeae, Secamonoideae) lacked any candidate PA ions in the PREC 120 (with MCA) scans (Table S1). Tribes Fockeeae, Eustegieae, and Ceropegieae were not sampled.

### 2.4. Wrightieae

Wrightieae are sister to the rest of the APSA clade and include three genera (Livshultz et al., 2007). Likely PAs were detected in leaves of 5 of 11 species of *Wrightia* R.Br. (Table 2) but none of the three species of *Pleioceras* Baill. and *Stephanostema* K.Schum sampled (Table S1). The most frequent candidate PA present yields an ion of m/z 432.5, containing both m/z 120 and 138 fragments. However, there is some evidence for coelution with an m/z 300.2 compound, as a low intensity m/z 300.2 ion was also associated with the TIC of the m/z 432.5 compound. Seeds of *Wrightia tinctoria* also contain an m/z 432.4 compound with fragmentation in both PREC 120 and PREC 138 (Table 2). Within the chromatography associated with the m/z 432.4 ion, m/z 270.2, 300.2 and 328.3 appeared with lower abundances (counts per second). Alterations to the chromatography method associated with these scans can increase the resolution of these compounds currently co-eluting within the chromatography, thereby increasing the sensitivity to these ions by diminishing ion suppression. Multiple other PREC 120 only ions appeared in these samples including m/z 210.2, 352.1, and 460.0. We suggest with low confidence these molecular ions are associated with PAs.

Five of the *Wrightia* species sampled here were included in a molecular phylogenetic analysis (Livshultz et al., 2007). The distribution of PAs on the *Wrightia* topology is homoplastic, but the potential utility of PAs as a chemotaxonomic character for the infra-generic classification of *Wrightia* warrants more thorough sampling.

Previously reported alkaloids from *Wrightia* species include steroidal (pregnane) alkaloids from *Wrightia pubescens* subsp. *laniti* (Blanco) Ngan (syn. *Wrightia javanica* A.DC.) (Kawamoto et al., 2003). This species is sampled here (Table S1) and determined as PA negative. Three species, *W. tinctoria*, *W. dubia* Spreng., and *W. arborea* (Dennst.) Mabb. (syn. *W. tomentosa* Roem. & Schult.), are used as sources of indigo (Ngan, 1965), and the indole-containing compounds indigotin and indirubin are known from *W. tinctoria*, as well as the quinolizidine, tryptanthrin (Khyade, 2014). Two of these three species, *W. tinctoria* and *W. dubia*, are here reported as likely containing PAs (Table 2) while the third, *W. arborea*, is reported as PA negative (Table S1).

### 2.5. Malouetieae

Eleven species from nine of 13 genera of Malouetieae were sampled. PAs were detected with high confidence in *Eucorymbia alba* Stapf (two samples) and *Galactophora schomburgkiana* Woodson (one sample) (Table 2). This is the first report of secondary metabolites from these genera. No PAs were detected in the single sample of *Galactophora crassifolia* (Müll.Arg.) Woodson analyzed (Table S1). Evenly massed ions were detected in the PREC 120 (with MCA) scan in one of two samples of *Holarrhena curtisii* King & Gamble (m/z 284.3, 300.1) (Table S1). *Kibatalia macrophylla* (Pierre) Woodson (one of two samples) contained three possible PAs in its PREC 120 (with MCA) mass spectrum, m/z 284.3, 302.4, and 314.0 (Table S1).

The only previous report of PAs in Malouetieae used the Mattocks’ test (Mattocks, 1967), a colorimetric assay with absorption at 565 nm, to detect unsaturated PAs in the seeds of *Holarrhena pubescens* (Arseculeratne et al., 1981). In our study, crude extracts of *Holarrhena pubsescens* leaves were evaluated for PAs via LC-MS/MS, with no strong evidence of PAs in the PREC scans. In seeds of *H. pubescens*, two even-massed ions were identified in the PREC 120 scan only, m/z 476.4 and 498.4 (Table 2). These are considered candidate PAs with low confidence. The specialized metabolites of *H. pubescens* (syn. *H. antidysenterica* (L.) Wall. ex A.DC., *H. floribunda* T.Durand & Schinz) are well studied, with steroidal alkaloids and glycosides of primary interest for their potential bioactivity (Kumar et al., 2007; Sinha et al., 2013). Three steroidal amino-glycosides previously reported from *H. pubescens* (holantosine B, D, and F, MW=475.3 Da) correspond in mass to one of the ions detected in PREC 120 (Afendi et al., 2011, queried 16.07.2020; Janot et al., 1970). Analysis of standards and/or alternative analytical techniques would be required for absolute identification of the compounds detected here.

### 2.6. Echiteae, Odontadenieae, Mesechiteae

These three tribes form a well-supported monophyletic clade of predominantly New World genera (Fishbein et al., 2018), but the boundaries among them have been redrawn several times (Morales et al., 2017; Simões et al., 2004). Morales et al. (2017) proposed that presence of PAs is a useful chemotaxonomic character for delimiting Echiteae from Odontadenieae and identified their absence in *Pentalinon* Voigt as supportive of its transfer to Odontadenieae. We sampled eight of fourteen Echiteae, five of nine Odontadenieae, and three of six Mesechiteae genera. PAs were detected in seven of eight sampled Echiteae genera. Only the sampled *Prestonia* species (*P. robusta* Rusby, *P. tomentosa* R.Br.) had no evidence of PAs in the PREC 120 (MCA on) scans (Table S1), although PAs have been previously reported from *P. amabilis* J.F.Morales, *P. quinquangularis* Spreng. [syn. *P. acutifolia* (Benth. ex Müll.Arg) K.Schum], and *P. portobellensis* Woodson (Burzynski et al., 2015). PAs in *Echites* and *Parsonsia* have been previously reported (Burzynski et al., 2015), but PAs in *Bahiella* J.F.Morales, *Macropharynx* (syn. *Peltastes*), *Temnadenia*, *Rhodocalyx* Müll.Arg., and *Laubertia* A.DC. are here reported for the first time, although Brown (1987) had previously suggested their presence in *Macropharynx* and *Temnadenia* based on indirect evidence. Exemplars of these genera had molecular ions appearing at consistent retention times in PREC 120, 138, and 156 scans (Table 2), fragmentation patterns characteristic of the cyclic triesters known from the positive control, *Parsonsia alboflavescens* (Table 1). Among the 22 species sampled from other two tribes (Table S1), only one, *Mandevilla boliviensis* (J.J.Veitch) Woodson (Mesechiteae), yielded an ion detectable in the PREC 120 scan, but not in PREC 138 or 156 (Table 2). We have low confidence in the identification of this compound as a PA, but this result requires further investigation. Overall, this pattern supports (Morales et al., 2017) hypothesis of the chemotaxonomic utility of PA presence for circumscription of Echiteae.

PAs are present in all Echiteae genera studied, but presence of detectable PAs varies both among samples of congeneric species and of conspecific individuals (Table S1, Table 2). Also, the diversity of PAs within a sample varied greatly from 18 candidate compounds in *Parsonsia alboflavescens* (Table 1) to one in *Temnadenia odorifera* (Vell.) J.F.Morales (Table 2). We have evidence that some of this variation is due to plasticity (see Section 2.11), but the role of genetics remains uncertain. Extensive sampling of species and individuals within Echiteae genera will be necessary to determine if there are any species-specific patterns useful for delimiting sub-tribal or infra-generic taxa.

### 2.7. Apocyneae

We sampled 36 species from 17 of 21 genera of Apocyneae. In this tribe, PAs had been previously reported from *Anodendron affine* Druce (Sasaki and Hirata, 1970) and *Amphineurion marginatum* (Roxb.) D.J.Middleton (Colegate et al., 2016). These species belong to two early diverging lineages of Apocyneae (subtribes Papuechitinae and Amphineurinae, respectively) and Livshultz et al. (2018b) suggested that PAs were potentially a chemotaxonomic character for delimiting these early diverging lineages from the rest of the radiation.

No PAs were detected with high confidence in any sample of Apocyneae. Two of three accessions of *Amphineurion marginatum* leaves possessed a molecular ion of m/z 320 which appeared only in the PREC 120 scan with associated chromatography at a late retention time of ~15 mins (>3 to 1 S/N) (Table 2). Fragmentation of a PA with an m/z 320 via PIS by Colegate et al. (2016) was consistent with our PREC 120 findings, as that compound was found to only result in m/z 120 fragment with no m/z 138 or 156 fragments. HRMS was utilized to propose a molecular formula for the m/z 320 ion, C_18_H_26_NO_4_, and therefore it was tentatively identified as an open-chain diester (Colegate et al., 2016). Colegate et al. (2016) reported that leaves of *Amphineurion marginatum* contained the lowest percentage by dry weight of PAs (0.02%) compared to roots (0.13%) and stems (0.09%). This may account for the lack of PA structural diversity and abundance observed in our leaf samples but other factors, environmentally induced plasticity, genetic variation, cannot be ruled out in the present study. A product scan and/ or HRMS would need to be completed on the m/z 320 molecular ion in our *Amphineurion marginatum* leaf samples to fully understand the fragmentation pattern and identity of this compound.

The unusual PAs previously identified in *Anodendron affine* have a platynecine core (Sasaki and Hirata, 1970) that would not be detectable with our strategy. Thus the occurrence and distribution of platynecine-derived PAs in other Apocyneae species also remains an open question.

### 2.8. Marsdenieae

We sampled 26 species from 10 of 27 genera of tribe Marsdenieae. PAs were identified with high confidence in samples of *Marsdenia tinctoria* R.Br. and *Marsdenia glabra* Constantin (Table 2). The current taxonomic concept of *Marsdenia* R.Br. is highly polyphyletic (Rodda et al., 2020), and the segregation of monophyletic genera from *Marsdenia sensu lato* is ongoing (Espírito Santo et al., 2019). *Marsdenia tinctoria* (the type species) and *M. glabra* belong to the small group of species segregated as *Marsdenia sensu stricto* (Bullock, 1956; Forster, 1995). Presence of PAs may be a useful chemotaxonomic character for recognition of *Marsdenia sensu stricto*, pending further sampling. The most frequently reported natural products from species of *Marsdenia sensu lato* and other genera of Marsdenieae are steroids, steroidal alkaloids, and steroidal glycosides including the antisweet gymnemic acids from the medicinal plant *Gymnema sylvestre* (Retz.) R.Br. ex Sm. (Liu et al., 1992). All three classes of compounds have been previously reported from *Marsdenia tinctoria* (Chowdhury et al., 1994; Gao et al., 2009).

### 2.9. Asclepiadeae and Periplocoideae

We sampled 60 species from 25 of 105 genera of Asclepiadeae and 8 species from 6 of 33 genera of Periplocoideae. We identified compounds that might be PAs with low confidence in five (5) species of the closely related Asclepiadeae subtribes Metastelmatinae and Oxypetalinae (*Seutera angustifolia* (Pers.) Fishbein & W.D.Stevens, *Ditassa eximia* Decne., *Ditassa lenheirensis* Silveira, *Metastelma harleyi* Fontella, *Minaria magisteriana (Rapini) T.U.P.Konno & Rapini*) and in one of two samples of *Petopentia natalensis* (Schltr.) Bullock (Periplocoideae) (Table 2). The re-circumscription of genera in these species-rich Asclepiadeae subtribes is ongoing (Liede-Schumann et al., 2014; Liede-Schumann et al., 2005; Silva et al., 2012) and a chemotaxonomic approach has not been attempted. The compounds discovered in the present study, once they are identified, can form a nucleus for this research. Asclepiadeae as a whole are known for diverse steroidal compounds, including cardenolides (Endress et al., 2018). Steroidal alkaloids are widely distributed (Lee et al., 2000) and phenanthroindolizidine alkaloids are diagnostic of *Vincetoxicum* Wolf (Endress et al., 2018; Liede-Schumann et al., 2016; Liede, 1996; Staerk et al., 2005).

There are no previous reports of specialized metabolites from *Petopentia natalensis*, but a diversity of compounds have been reported from Periplocoideae including steroidal glycosides, cardenolides, steroidal alkaloids and, rarely, indoloquinoline alkaloids from species of *Cryptolepis* R.Br. (Endress et al., 2018; Paulo and Houghton, 2003; Tackie et al., 1991).

Rarely do retronecine-type PAs present without other diagnostic fragment ions such as m/z 138 and/ or 156 fragments. However, a few other PAs have been documented to contain a base peak of m/z 120 without possession of other key fragments. (Jeong and Lim, 2019) noted the tentative identification of two open chained retronecine-type diesters, symviridine and uplandicine, in which a base peak was *m/z* 120 and no other common fragments (m/z 138 or 156) were obtained in PIS scans at the collision energies utilized. Another open-chain diester, 7-acetyllycopsamine, was shown to fragment to a base peak of *m/z* 120 (Colegate et al., 2005; Jeong and Lim, 2019). It is yet to be determined if a common structural feature causes a tendency for particular PAs to result only in a *m/z* 120 fragment, but it does appear open-chained diesters preferentially result in observation of only an *m/z* 120 fragment at particular collision energies. This fragmentation pattern can yield only low confidence identification of retronecine-type PAs and leaves substantial ambiguity about the true identity of compounds that present with this pattern.

### 2.10. PA distribution and homospermidine synthase amino acid motifs

The first step of PA biosynthesis is catalyzed by the enzyme encoded by homospermidine synthase (*hss*) which evolved 7 times independently among angiosperm lineages via duplication and subfunctionalization of deoxyhypusine synthase (*dhs*), a gene of primary metabolism (Livshultz et al., 2018a). Livshultz et al. (2018a) identified a characteristic amino acid motif in HSS (VXXXD) that evolved from the ancestral IXXXN DHS motif each time HSS function and PA-biosynthesis evolved. *In vitro* experiments showed that the DHS of *Ipomoea neei* (Spreng.) O’Donell (wild-type IXXXN) mutagenized to a VXXXD motif had less DHS and slightly more HSS function (Kaltenegger et al., 2013). In Apocynaceae, Livshultz et al. (2018a) placed the gene duplication that gave rise to the functionally characterized HSS of *Parsonsia alboflavescens* early in the diversification of the APSA clade but after the divergence of Wrightieae from the rest of the clade. They hypothesized that species with *hss*-like genes that encode the VXXXD motif have a functional HSS and can produce PAs while species with *hss*-like genes that encode the DHS-like IXXXN motif or an intermediate IXXXD motif have lost PA biosynthesis.

Fifteen (15) of the species of from which *hss*-like genes were sequenced are included in the present study (Table 3). Three species with the VXXXD motif are identified with high confidence as producing PAs, three with low confidence, and four have no detectable PAs. One sampled species with the IXXXN motif produces PAs (*Marsdenia glabra*) while the other (*Asclepias syriaca* L.) lacks PAs and has a pseudogenized *hss*-like gene which encodes the IXXXN motif when the ORF is restored (Livshultz et al., 2018a). All three sampled species with the IXXXD motif lack detectable PAs. The relationship between motif and presence of PAs is not dichotomous as predicted by Livshultz et al. (2018) (Table 3). Both species with VXXXD and IXXXN motifs in their HSS-like enzymes can produce PAs. More sequencing of *hss*-like genes from the species included in the present survey is required to determine if there is any correlation between the identified motifs and PA biosynthesis and if any other amino acid substitutions might be functionally important. More infra-specific sampling and scans for PAs with alternative cores (Fig. 1) are required to confirm that species reported here as PA negative truly lack PAs. No *hss*-like gene was found in the transcriptome of *Wrightia natalensis* Stapf (a PA negative species, Table S1). It remains to be determined if PA-producing *Wrightia* species (Table 2) have an *hss* locus in their genomes.

**Table 3.**
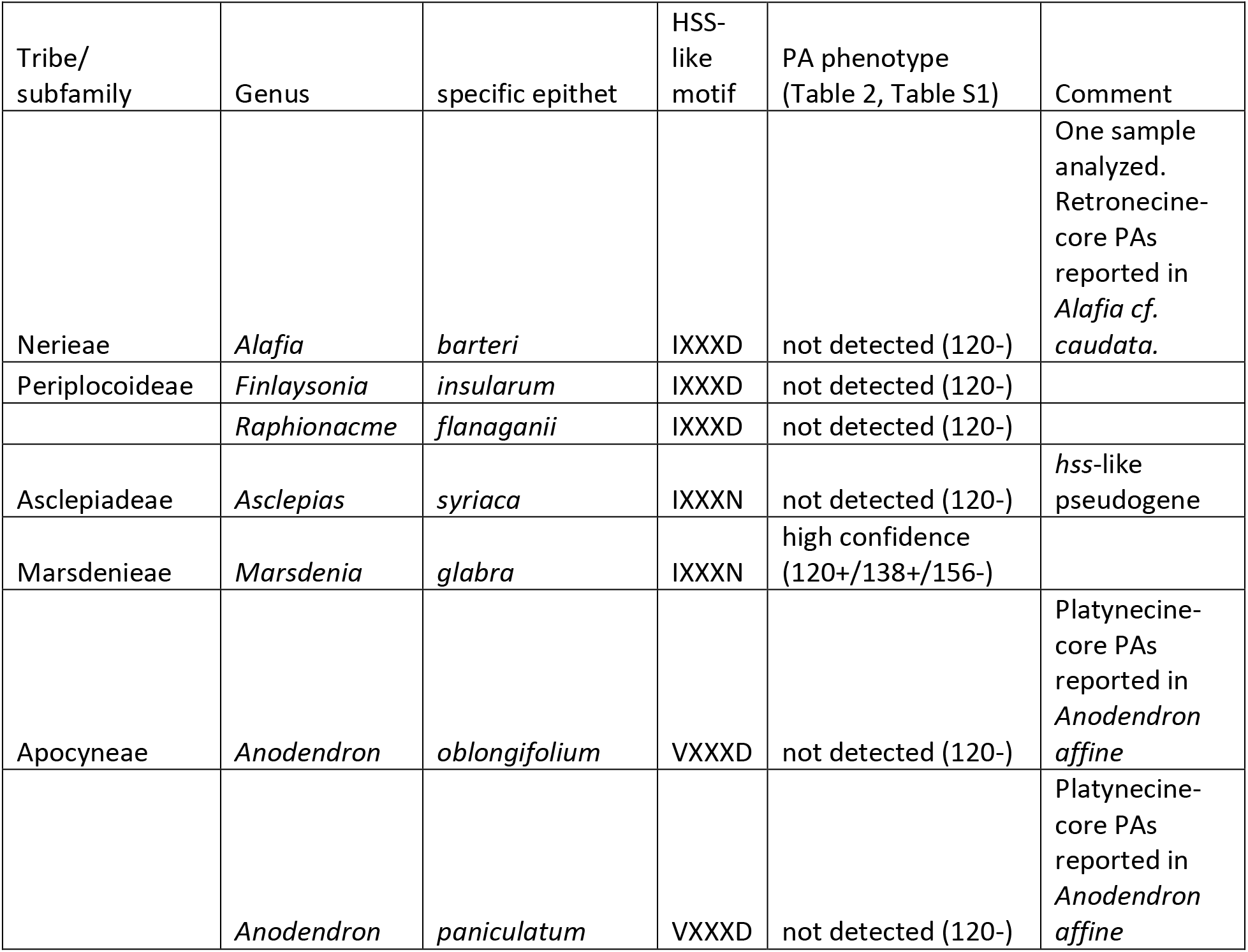
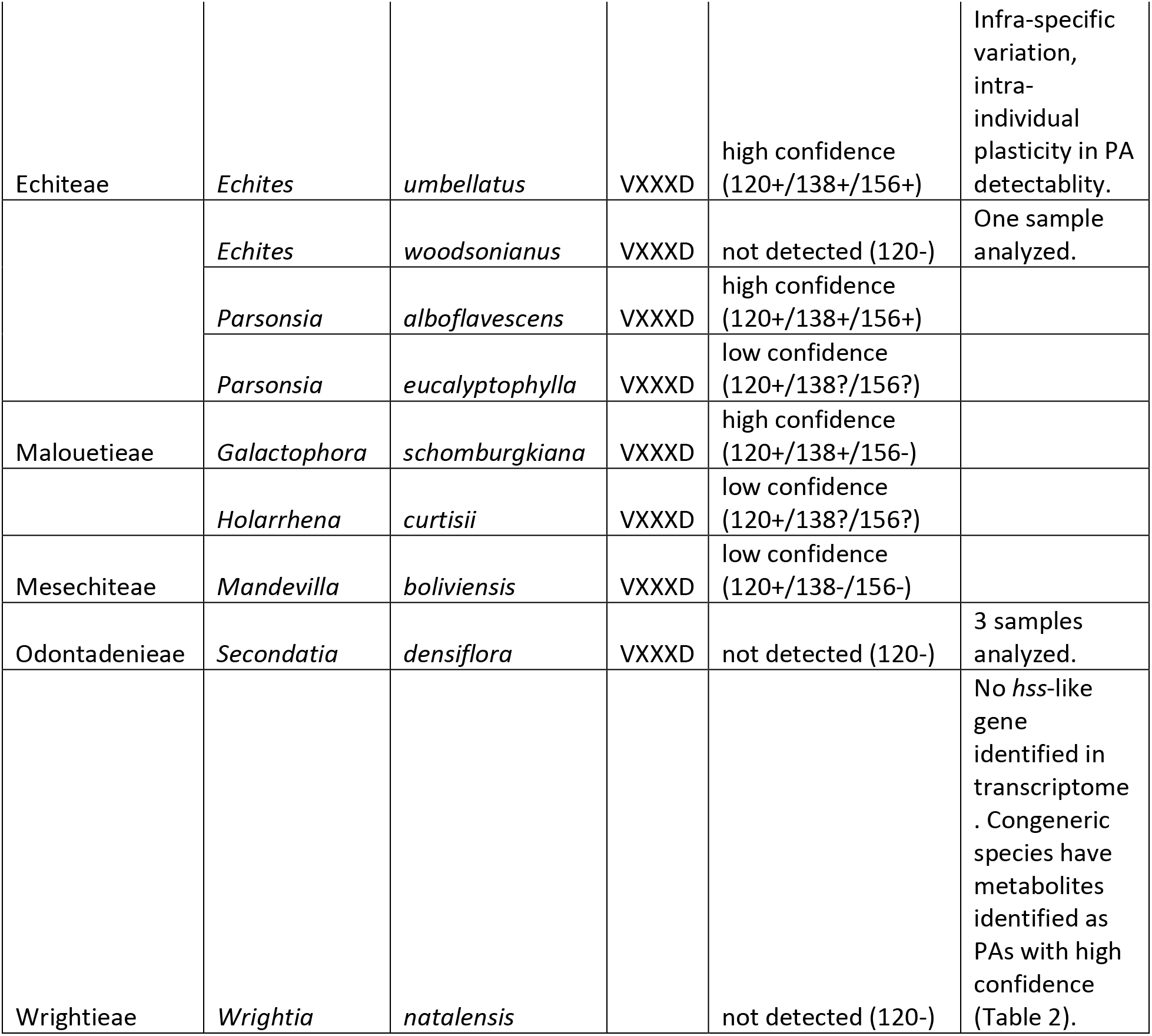
Amino acid motifs of *homospermidine synthase*-like genes (Livshultz et al., 2018a) from Apocynaceae species and PA phenotypes. The VXXXD motif is associated with evolution of a functional HSS in comparative analyses of PA-producing angiosperms.

### 2.11 Infraspecific variation in PA detectability

Tasca et al. (2018) were unable to detect PAs in a subset of *Echites umbellatus* herbarium specimens analyzed with a Q1 scan. Since some of the specimens analyzed in that study were very old, and because a PREC scan can detect PAs with greater specificity than a Q1 scan, we follow up here by analyzing leaf fragments from 19 relatively recent (1935-2006) herbarium specimens with PREC 120 (with MCA). Of these, nine (9) had detectable PAs and ten (10) did not (Table S2). We also evaluated 110 leaf samples from 96 cultivated plants in seven half-sib families established from wild collected fruits from Florida and the Bahamas (Table S3). Among all samples, 98 had PAs while 31 had no detectable PAs. PA detection was unrelated to extracted leaf mass. On average, less tissue was used for positive samples (mean=11.1 mg, SD=5.5 mg, N=98) than for negative samples (mean= 21.4 mg, SD=16.3 mg, N=31).

In five of the seven half sibling families analyzed, we detected PAs in the youngest leaves collected from all individuals sampled (9-14 plants/family). In two families, 5 of 12 and 7 of 19 individuals had no detectable PAs when first sampled (Table S3). The youngest leaves from fourteen (14) of the 31 individuals in these two families were re-sampled two years later. Among these plants, the youngest leaves of five (5) individuals maintained the same phenotype (two with detectable PAs, three with no detectable PAs), six (6) shifted from detectable PAs at time one to no detectable PAs at time two, and three (3) shifted from undetectable to detectable. Thus the variation in detectability of PAs observed among samples is, at least in part, due to phenotypic plasticity. Because all plants had already transitioned to flowering when they were sampled at time 1, and because the same approximate developmental stage was sampled at both time points (youngest leaves), environmental factors are implicated. However, the inconsistency in the direction of change, and the difficulty of exactly matching leaf developmental stages, means that ontogenetic variation is not completely excluded based on the current data. Genetic factors may also play a role in determining the presence of plasticity in two of our seed families but not in five others. More tightly controlled experiments are necessary to identify the causal factors.

Infraspecific variation is very commonly in distribution of specialized metabolites both among and within conspecific plants (Hartmann, 1996; Moore et al., 2014), included in those that produce PAs. For example, *Senecio vulgaris* L. showed distinction differences in specific PA concentrations depending on both developmental stage and season (Flade et al., 2019), with the total PAs present increasing with developmental stage.

### 2.12 Implications for human health

Apocynaceae species are frequently used medicinally and more rarely as food (Endress et al., 2018). Several of the species here reported with high confidence as producing PAs are consumed by people. Notably, commercially available seeds of *Wrightia tinctoria* showed evidence of compounds that are identified with high confidence as PAs (Table 2). Seeds of *Wrightia tinctoria* are consumed in Ayruveda and other medicinal traditions of India, both intentionally (Khyade, 2014), and as contaminants of *Holarrhena pubescens* seeds (Khan, 1987). We found contamination by *Wrightia tinctoria* in our purchased package of *Holarrhena pubescens* seeds. Other organs of *Wrightia tinctoria* (leaves, bark) are also consumed (Reddy et al., 1999), as is *Wrightia antidysenterica* (L.) R.Br. (syn. *Walidda antidysenterica* (L.) Pichon) (Wickramaratne et al., 2015), another species identified with high confidence as PA positive (Table 2). On the other hand, we assign low probability to presence of retronecine-type PAs in seeds of *Holarrhena pubescens*, separated from contaminating *Wrightia tinctoria* seeds. Because specialized metabolites of *Holarrhena pubescens* are so well studied (Sinha et al., 2013) and because neither of the candidate compounds fragmented in the PREC 138 and 156 scans (Table 2), we consider it more likely that the m/z 476 and 498 ions identified in PREC 120 correspond to other primary or secondary metabolites. We suggest that the Arseculeratne et al. (1981) report of PAs in *H. pubescens* seeds derives from either non-specificity of their colorimetric assay or contamination by *Wrightia* spp. But no final conclusion can be reached until the low-confidence compounds that we found are definitively identified.

*Marsdenia tinctoria* is not as readily available in the global marketplace as *Wrightia tinctoria* and *Wrightia*-contaminated *Holarrhena pubescens*, but *Marsdenia tinctoria* is consumed as a remedy in the traditional Chinese medicine system and elsewhere in southeast Asia (Chowdhury et al., 1994; Yan-Jiao, 2013). Any potential risk to people from consumption of *Wrightia* and *Marsdenia* products is unknown at this point. Further research, beginning with NMR characterization of the compounds we identified (Table 2), is warranted.

Flowers of *Echites panduratus* A.DC. (syn. *Fernaldia pandurata* Woodson, *Urechites karwinskii* Müll.Arg.) are consumed as a pickled condiment, “loroco,” in the cuisines of several Central American countries, and young shoots are eaten as a green vegetable (Morton et al., 1990). A keto dihydro-PA, loroquine, was reported from the roots of *Echites panduratus*, but was not detectable in the shoots (del Castillo et al., 1970). Morton et al. (1990) also reported absence of indole alkaloids in aerial plant organs. Burzynski et al. (2015) did not detect PAs in sampled leaves, but Colegate et al. (2016) commented on unpublished evidence of retronecine-type PA presence in roots. In the present study, we re-analyzed the same specimen sampled by Burzynski et al. (2015) and confirmed absence of detectable PAs in PREC 120 (Table S1) and in a second sample from another plant, confirming no detectable compounds in PREC120/138/156 (Table 2). We also sampled shoot tips, young and mature leaves, and roots from a third plant, and found evidence of PAs with PREC 120 in mature leaves and roots (Table S1). These results need follow up with PREC 138/156 scans for confirmation. Assessment of any human risk from PAs related to consumption of *Echites panduratus* must begin with the identification and quantification of PAs in preparations as consumed. Broad sampling is necessary, given the potential for ontogenetic, environmentally-induced, and genetic variation.

### 2.13 Limitations of the current approach: sampling strategy

The broad taxonomic sampling that we sought to accomplish in this study, means that most species were represented by only one or a few samples. The variation in PA detectability that we documented in *Echites umbellatus* (ca. 50% of herbarium specimens with no detectable PAs, Table S2), shows that there is a high probability that some of the species here reported as lacking retronecine-type PAs in the leaves, will be reported to have them when more samples are analyzed. Our sampling is overwhelming biased toward young leaves, since most of the samples were acquired for DNA extraction. PA accumulation often has a distinctly organ and developmental stage specific pattern that is variable among species (Hartmann and Zimmer, 1986; Irmer et al., 2015; Stegemann et al., 2018). In *Amphineurion marginatum* and *Alafia cf. caudata* (Apocynaceae), PA concentration and diversity was highest in root and lowest in leaves (Colegate et al., 2016). In *Echites panduratu*s, we detected PAs in old but not in young leaves (Table S1). If these patterns are consistent across Apocynaceae species, our study is biased toward underreporting of PA occurrence.

### 2.14. Limitations of the current analytical approach: alternative necine cores and N-oxides

The current MS methods exclude the possibly of identifying alternative necine bases, namely otonecine and playtnecine. Identifying these bases would require additionally MS scans to target the diagnostic fragments specific to these structures. Otonecine-type PAs, for example clivorine, readily exhibit m/z 122, 150, and 168 fragments; with m/z 150 and 168 ions being most diagnostic, similar to the m/z 120 and 138 diagnostic ion fragments associated with retronecine-type PAs (Lin et al., 1998). Alternatively, saturated necine bases of playtnecine-type PAs, have a diagnostic fragment ion of m/z 140 and a characterization ion of m/z 122 (Lin et al., 1998). Since alternative bases were not pursued, the full diversity of PAs present in our samples cannot be determined with the current methodologies.

Additionally, the N-oxide form of PAs is often more prevalent in plants than the free base, likely due to the higher solubility and lower toxicity of the N-oxide form relative to the free base form of PAs (Hartmann and Witte, 1995; Hartmann and Zimmer, 1986). Free base PAs can be differentiated from their N-oxides via *in situ* reduction, typically via zinc dust (Colegate et al., 2005; Colegate et al., 2016; Nishida et al., 1991). Many examples of the prevalence of N-oxides in Apocynaceae have been documented. In *Alafia cf. caudata,* 10 of 15 PAs isolated in roots were N-oxides (Colegate et al., 2016). In *Parsonia straminea* (R.Br.) F.Muell., 96% of PAs identified via high resolution accurate mass (HRAM) LC-MS/MS in leaves and stems were N-oxides (Hungerford et al., 2019). Therefore, not accounting for the presence of N-oxides in samples can lead to the possibility of underrepresenting the amount of PAs present, in addition to falsely characterizing some species as non-PA producing.

N-oxides can be detected without derivatization via MS, resulting in unique diagnostic fragments in addition to those produced by the free base form of PAs. Typically, N-oxide diesters (open and closed) present with m/z 120 and 138 ions, in addition to m/z 136 and 118 fragments (Ruan et al., 2012; These et al., 2013). A PIS (product scan) of lycopsamine N-oxide, under the instrumental parameters utilized, resulted in one ion pair, m/z 138-136 (Ruan et al., 2012). However, a high intensity m/z 172 fragment was observed, representative of the m/z 156 fragment typical of open-chain monoesters with an associated oxygen atom. These et al. (2013) alternatively observed both ion pairs (m/z 118-120 and m/z 136-138) and the m/z 172 fragment when lycopsamine N-oxide was subject to PIS. Fragmentation of the retronecine core was frequently observed in product scans of N-oxides under the collision energies utilized, with m/z 94 the most prominent ion (Ruan et al., 2012; These et al., 2013).

The PREC scans employed in this study were designed to specifically target fragments inherent to pro-toxic, free base, 1,2-unsaturated retronecine-type PAs. Standards of this type were used to ensure proper fragmentation of these PAs. Burzynski et al. (2015) utilized PREC and MRM scans to identify the m/z 120 fragment in masses associated with the N-oxide of parsonsine-type PAs. Our current PREC scans were also able to detect masses associated with tentatively identified N-oxide structures determined by Burzynski et al. (2015) (Table 1). This indicates our methods have some specificity for N-oxides possibly because the N-oxide and free base form of retronecine-type Pas typically present with m/z 120 and 138 fragments. However, more specific structural features must be targeted in our MS protocols to increase the specificity for detection of N-oxide PAs. This is particularly important since (Avula et al., 2015; Avula, 2015) and Colegate et al. (2016) note detection of N-oxide PAs through PIS (utilizing their respective experimental MS parameters) that display either only m/z 120 or m/z 138 fragments or neither fragment. With our current method, the m/z 120 fragment had to be present in order for identification of a retronecine-type PA to be considered.

## 3. Conclusion

We have conducted the first systematic survey of Apocynaceae for retronecine-type PAs using high throughput LC-MS methods, PREC 120 (with MCA) and PREC 120/ 138/ 156. We confirm that PAs have a scattered distribution within one lineage of the family, the APSA clade, as predicted from previous reports in the phytochemical literature and from the distribution of *homospermidine synthase*-like genes, which encode the first enzyme of the PA biosynthetic pathway. PAs are confirmed as a useful chemotaxonomic character for delimiting Echiteae (PAs present in at least one species of all genera tested to date) from the closely related Odontadenieae and Mesechiteae (only one species yielded an ion identified with low confidence as a potential PA). In contrast to Echiteae, only a few of the tested species from other previously reported PA positive tribes (Nerieae, Malouetieae, Apocyneae) have detectable PAs. We report likely PAs for the first time in Wrightieae (species of *Wrightia*) and Marsdenieae (species of *Marsdenia sensu stricto*). Discovery of these novel PA sources has potential implications for human health since two of the species that we identify as likely containing PAs, *Wrightia tinctoria* and *Marsdenia tinctoria*, are used medicinally. We find clear evidence of infra-specific variation in PA-detectability in young leaves of *Echites umbellatus* and confirm that phenotypic plasticity is one of the causes. Thus it is worth following up with further sampling of genera and species that are reported here as lacking detectable PAs but that are closely related to PA-positive taxa. Targeted studies of organ-specific, developmental stage, environmental, and genetic determinants of the PA phenotype are also necessary. The more complete picture of the taxonomic distribution of PAs in Apocynaceae provided by this report will facilitate research into the evolution of the PA biosynthetic pathway and into co-evolution of Apocynaceae and their PA-adapted herbivores.

## 4. Experimental

### 4.1. Plant materials

Most tissue samples are young leaves, collected into desiccant from the living plant, and stored at room temperature. A minority of samples are young leaves removed from herbarium specimens. Most of these specimens had been dried at ca. 30 °C when collected from the living plant, potentially with alcohol and/or mercuric chloride treatment, and stored at room temperature (Tasca et al., 2018). Shoot tips, young and mature leaves, and young roots collected from a cultivated accession of *Echites panduratus* were dried at room temperature. Seeds of *Holarrhena pubescens* and *Wrightia tinctoria* were purchased from retail outlets in the United States and stored at room temperature. Seeds were identified based on morphology, (Khan, 1987) and sorted under magnification to produce pure samples, free of admixture, for analysis. Each analyzed tissue sample is vouchered by a specimen deposited in a publicly accessible herbarium (Table S1). Taxonomy follows Endress et al. (2018) as updates by Espírito Santo et al. (2019) and Rodda et al. (2020).

Half-sib families of *Echites umbellatus* were established in fall 2013, spring 2014, and fall 2014, by planting the seeds from individual wild-collected fruits and cultivated in the greenhouse of Florida International University. Seeds were started in small-celled seedling trays, then transplanted into Metromix in 4” pots, and finally into 3 gallon pots, with three bamboo poles attached at the top to support the vines. Plants were grown under ambient light with summer high temperatures usually ca. 29 – 35 °C with ca. 75-98% humidity. The youngest leaves from each plant were collected from April-June 2016. The youngest leaves from a subset of these plants were re-sampled in March-April 2018. Leaves were collected immediately into desiccant and stored at room temperature. All plants that were sampled twice had already flowered before the first sampling in 2016.

#### 4.1.2. Chemicals and PA Standards

PA standards (heliotrine, jacobine, retrorsine, and europine) were purchased from Phytolab GMBH (Vestenbergsgreuth, Germany). Lycopsamine and monocrotaline were purchased from Toronto Research Chemicals (Toronto, Ontario, Canada). Standards were used as received and dissolved in HPLC-grade methanol prior to use. Acetonitrile and water for HPLC use were also HPLC-grade.

### 4.2. Extraction and isolation- Dried Leaves

The extraction method utilized in this study follows Tasca et al. (2018) to maximize PA recovery. Leaves were placed in 1.7 mL Eppendorf tubes and manually crushed with a microspatula for 15 seconds. Approximately 4-60 mg of dried leaf tissue was used for each extraction. Each specimen was then suspended in 750 μL of methanol, vortexed for 5 seconds, left to rest for 24 hours, and vortexed again for 5 seconds. The crude extracts were then removed and transferred to a second 1.7 mL Eppendorf tube, followed by 2 × 250 μL methanol washes. The methanol was then evaporated from the crude extracts *in vacuo* in a DR 120 SpeedVac with the drying rate on low. The crude extracts were reconstituted in 80 μL of methanol and vortexed for 30 seconds. The reconstituted samples were then transferred to 0.25 mL plastic autosampler vials with 1 mL Norm-Ject® disposable syringes fitted with Acrodisc CR 13 mm, 0.2 μm PTFE membrane filters.

#### 4.2.1. Extraction and isolation-from seeds

To maintain congruency in sample preparation between leaves and seeds; seeds were also extracted according to methods outlined in Tasca et al. (2018) with a few modifications. Between 250-500 mg of seeds were ground with a mortar and pestle until a fine powder was achieved. Ground seeds were then transferred to a 10 ml conical tube and 3 ml of HPLC-grade methanol was added. Samples were then sonicated (not exceeding 60°C) for 12 hours. The sample was then vortexed for 30 seconds and subsequently centrifuged for 10 minutes. The supernatant was then removed and placed into a 1.7 mL Eppendorf tube to be evaporated *in vacuo* on a DR 120 SpeedVac with the drying rate on low. Subsequently, the pellet was washed with another 3 ml of HPLC-grade MeOH. The resuspended pellet was vortexed for 30 seconds, centrifuged for 10 minutes, with the supernatant again dried on the SpeedVac. The dried material in the Eppendorf tube was then resuspended in 250 μL of methanol and vortexed for 30 seconds. The reconstituted sample was then transferred to a 0.25 mL plastic autosampler vial with a 1 mL Norm-Ject^®^ disposable syringe after being passed through an Acrodisc CR 13 mm, 0.2 μm PTFE membrane filter.

### 4.3. Liquid Chromatography Methods

All samples, regardless of the fragment ion being targeted in the PREC scan, were run on the same liquid chromatography method to allow for comparison of retention times across scans, although retention times were uncorrected. Crude extracts were separated via reverse-phase HPLC (Shimadzu, LC-20AB, Kyoto, Japan) with an ACE C18 column (3 μm particle, 150 × 4.6 mm) fitted with an ACE C18 guard column secured with a guard column holder (Advanced Chromatography Technologies Ltd., Aberdeen, Scotland). A binary mobile phase consisting of acidified water (0.1 % formic acid, v/v; Solvent A) and acidified acetonitrile (0.1 % formic acid, v/v; Solvent B) was utilized at a flow rate of 0.5 mL/min. Chromatography lasted 22 minutes for each run with the following gradient: 10 % B (2 min hold) ramped to a mobile phase concentration of 75 % B over 15 minutes, ramped to 95 % B (3 min hold), then 10 % B for the remainder of the run. 10 μL of sample was injected for each run.

To ensure the reproducibility of the HPLC method, a column wash method was utilized every 10-12 samples. This method consisted of a 20-minute hold of solvent B. The frequent use of column washes successfully mitigated retention time shift as the leaf tissue matrix could alter the separation capacity of the column after multiple runs. Prior to separation of the crude extracts, PA standards were analyzed via PREC methods on the washed and equilibrated HPLC column to establish consistency of the chromatography.

### 4.4. Mass Spectrometry methods and PA positive identification

All mass spectra were collected using an Applied Bio-Systems API 2000 triple quadrupole mass spectrometer (Waltham, Massachusetts) in positive electrospray ionization (ESI+) mode. Preliminary PREC 120 scans utilizing multiple channel acquisition (MCA) were utilized to tentatively identify PAs with a m/z 120 fragment. MS compound dependent parameters were kept the same for PREC 120 scans with and without MCA. Samples containing a molecular ion with a m/z 120 fragment were determined to be positive (with low confidence) when chromatography (greater than 3:1 S/N) present between 9-15 mins (40-75% ACN) in the total ion chromatogram (TIC; PREC 120 ion intensity vs. time) was associated with an evenly-massed molecular ion of intensity (cps; counts per second) greater than or equal to 1×10^4^. Select samples were then run on PREC 120/138/156 methods to elucidate additional fragments associated with even-massed molecular ions appearing in the mass spectrum of 120 MCA scans and to determine the possible presence of PAs in samples containing ambiguous chromatography associated with odd and even molecular ions in their mass spectrum.

Samples run on the three precursor scans specific to the fragments of the retronecine core were determined to be PA positive (with high confidence) if evenly massed molecular ions, greater than or equal to 1×10^4^ cps, were observed in the TIC with greater than 3:1 S/N between 9-15 minutes present within approximately 0.3 mins chromatographically in either the PREC120 and 138 scans or PREC 120, 138, and 156 scans.

The PREC scan for the m/z 120 fragment was optimized using a monocrotaline commercial standard with a declustering potential of 33 V and a collision energy of 45 V, producing a 120 *m/z* fragment in Q3. Alternatively, to establish the optimal declustering potential and collision energy with which to obtain the m/z 138 and 156 fragments, standards (monocrotaline, retrorsine, jacobine, heliotrine, europine, and lyposamine) were subject to product scans. Compound dependent parameters were manipulated to alter the intensity of the m/z 138 and 156 fragments. Ultimately, a declustering potential of 40 V and collision energy of 40 V was utilized for the PREC138 scan. The PREC156 scan alternatively employed a declustering potential and collision energy of 40 V and 50 V, respectively. All other MS parameters were kept the same across each of the PRECs scans with a scan mass range of m/z 200-500. Both system control and data analysis were performed on Analyst 1.6.2 software.

## Acknowledgements

This work was supported by the National Science Foundation, grants DEB-1655660 and DEB-1655663 to KM and TL. Mary E. Endress, David J. Middleton, Paul E. Forster, Cassia Bitencourt, Alessandro Rapini, and the herbarium of the Fairchild Tropical Garden (FTG) provided many samples and/or identified voucher specimens for this study. Bria Gillard and Mark Olson are thanked for their mass spectrometric suggestions and insights.

## APPENDIX A. Supplementary data

Table S1: Survey of Apocynaceae samples, voucher specimens and results of PREC 120 (MCA on) scans. Table S2: Survey of *Echites umbellatus* herbarium specimens, voucher specimens and results of PREC 120 (MCA on) scans. Table S3: *Echites umbellatus* half-sib families, voucher specimens and results of PREC 120 (MCA on) scans.

